# Discriminating predation attempt outcomes during natural foraging using the post-buzz pause in Japanese large-footed bat *Myotis macrodactylus*

**DOI:** 10.1101/2021.08.27.457998

**Authors:** Yuuka Mizuguchi, Emyo Fujioka, Olga Heim, Dai Fukui, Shizuko Hiryu

## Abstract

Bats emit a series of echolocation calls with an increasing repetition rate (the terminal buzz), when attempting to capture prey. This is often used as an acoustic indicator of prey-capture attempts. However, because it is directly linked to foraging efficiency, predation success is a more useful measure than predation attempts in ecological research. The characteristics of echolocation calls that consistently signify predation success across different situations have not been identified. Due to additional influencing factors, identification of these characteristics is particularly challenging for wild bats foraging in their natural environment compared to those in flight chambers. This study documented the natural foraging behavior of wild Japanese large-footed bat *Myotis macrodactylus* using synchronized acoustic and video recordings. From the video recordings, we could assign 137 attacks to three outcome categories: prey captured (51.8%), prey dropped (29.2%), and failed attempt (19%). Based on previous indications from laboratory studies that the length of the silent interval following the terminal buzz (post-buzz pause) might reflect the prey capture outcome, we compared post-buzz pause durations among categories of attack outcomes. The post-buzz pause was longest in the case of successful capture, suggesting that the length of the post-buzz pause is a useful acoustic indicator of predation success during natural foraging in *M. macrodactylus*. Our finding will advance the study of bat foraging behavior using acoustic data, including estimations of foraging efficiency and analyses of feeding habitat quality.

**Summary statement:** We investigated the natural foraging behavior of wild *Myotis macrodactylus* and found that the length of the post-buzz pause is a useful acoustic indicator of predation success.

## Introduction

Echolocating bats obtain information about the outside world by emitting sonar signals and listening to their echoes. This information helps them to find and capture prey, avoid obstacles and navigate (Fenton, 1990; Simmons et al., 1979). During foraging, insectivorous bats emit a series of echolocation calls that vary in pulse length, frequency structure, and repetition rates, and can therefore be grouped into three phases: the search, approach, and terminal phases (Griffin et al., 1960; Kalko and Schnitzler, 1993; Schnitzler and Kalko, 2001; Simmons et al., 1979). The “terminal buzz” is a rapid increase in the repetition rate that occurs just before a capture (Fig. 1A, Fujioka et al., 2014; Schumm et al., 1991). This unique acoustic feature reflects a “fast decision response” (Geberl et al., 2015), and has been used as an important acoustical indicator of capture attempts by bats during foraging (Britton and Jones, 1999; Griffin, 1958; Jakobsen and Surlykke, 2010; Racey and Swift, 1985).

**Figure 1.**
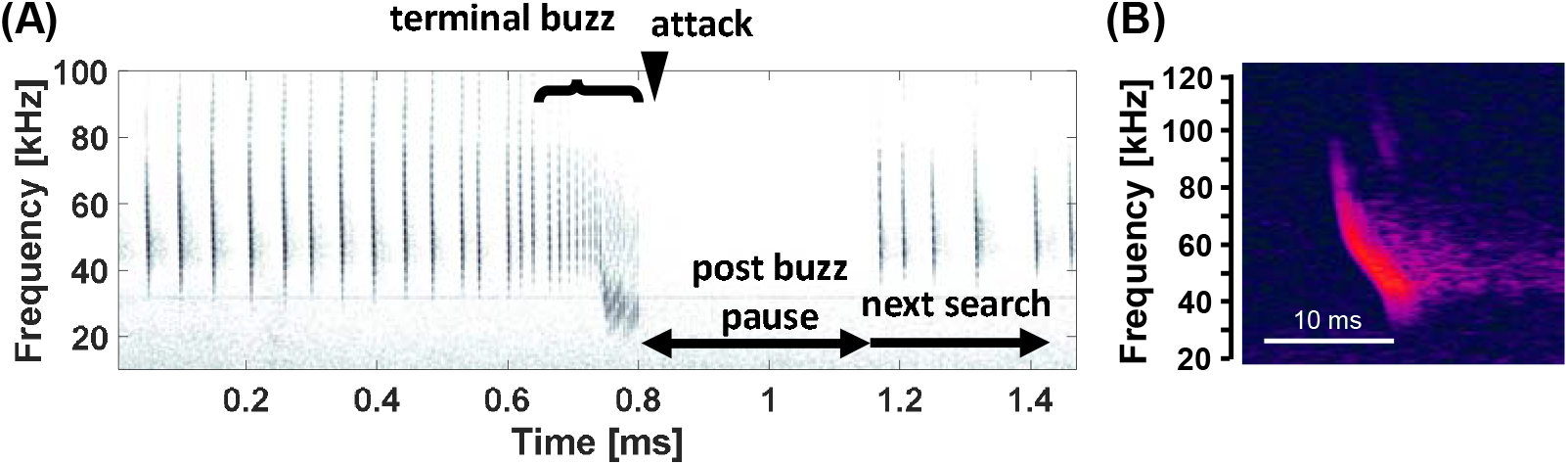
(A) Spectrogram of an echolocation call sequence from *Myotis macrodactylus* during prey capture. The terminal buzz is emitted just before attacking the prey. The time between the attack and the next search pulse is called the post-buzz pause. (B) Example search-phase echolocation call from *M. macrodactylus*. The fundamental frequency of this frequency-modulated (FM) pulse drops from about 90 to 40 kHz, with a peak frequency of 50 kHz.

Bats do not always succeed in capturing prey. Visual observations, photographs, and video recordings of foraging bats have shown that they sometimes drop prey or fail to capture it (Acharya and Fenton, 1992; Britton and Jones, 1999; Kalko, 1995; Schnitzler et al., 1994). For instance, *Eptesicus nilssonii*, an aerial-hawking bat, was found to successfully capture moths during natural foraging in only 35–36% of attempts (Rydell, 1992; Rydell, 1998). The capture success of the pipistrelle bat varies according to prey size, i.e., 60–70% of small-sized insects and 30–40% of large-sized insects, such as moths, were captured successfully (Kalko, 1995). Photographs of *Myotis daubentonii* foraging above the water surface have shown that the bats sometimes miss their prey or mistake it for other objects, such as floating leaves (Kalko and Schnitzler, 1989). Therefore, acoustic measures of predation success are desirable. These could be used to estimate the foraging status of bats from their emitted echolocation sounds, enabling measurement of temporal changes in energy intake and bat foraging efficiency. The use of acoustic parameters to determine predation outcome would greatly contribute to our understanding of natural foraging behavior, with bats as a model.

Laboratory studies investigating the acoustic characteristics of predation success have found that the length of the silent period at the end of a terminal buzz (post-buzz pause) is longer when predation is successful compared to when it is unsuccessful (Acharya and Fenton, 1992; Britton and Jones, 1999; Surlykke et al., 2003; Übernickel et al., 2013). It is assumed that the length of the post-buzz pause reflects the time required for a bat to bend its head towards its tail membrane pouch and grasp the prey (Acharya and Fenton, 1992; Kalko and Schnitzler, 1989). Furthermore, if a predation attempt fails, the next search is expected to start earlier, leading to a shorter post-buzz pause. However, this relationship has not been confirmed in field experiments (Britton and Jones, 1999). Thus, to the best of our knowledge, the echolocation call characteristics that enable discrimination between successful and unsuccessful predation in naturally foraging bats have not yet been identified.

The purpose of this study was to investigate whether the length of the post-buzz pause could be used as an acoustic indicator of successful predation in bats during natural foraging. We hypothesized that the length of the post-buzz pause would depend on the attack outcome. In particular, if a bat successfully captured its prey, we expected to find a longer post-buzz pause, with a shorter pause reflecting a failed attempted, as demonstrated in a previous laboratory experiment with *Myotis daubentonii* (Britton and Jones 1999a). Furthermore, if a bat dropped its prey, we expected the post-buzz pause to have an intermediate length.

## Materials and Methods

### Study species and setting

The target species was Japanese large-footed bat *Myotis macrodactylus*, a member of the family Vespertilionidae. This species emit a frequency-modulated (FM) pulse with a fundamental frequency falling from approximately 90 to 40 kHz, and a second harmonic (Fig. 1B, Fukui et al., 2004; Luo et al., 2012). In Japan, these bats feed mainly on prey from the orders of Diptera, Trichoptera, and Lepidoptera (Funakoshi and Takeda, 1998), and typically trawl for prey by flying above the water surface (Luo et al., 2012). Bat species that predominantly use trawling for prey capture are particularly suitable for detailed observations of natural foraging behavior via acoustic and video recordings because the feeding sites can be easily identified.

Our study site was a 20 m diameter pond (Fig. 2A) in Tomakomai Experimental Forest (42°43’N, 141°36’E), a research facility of Hokkaido University in Tomakomai, Hokkaido, Japan. The pond is part of a Horonai stream (about 3 m wide) that enters the pond from one side and exits it on the other side (Fig. 2A). Bats regularly use the open space above this pond for foraging after sunset during early summer and fall. In most of the cases that we observed, the bats appeared upstream or downstream of the stream, foraged above the pond for a certain period of time, and exited via the downstream side of the stream. The average temperature, average humidity, weather conditions, and sunset times during the recording period are shown in Table 1.

**Figure 2.**
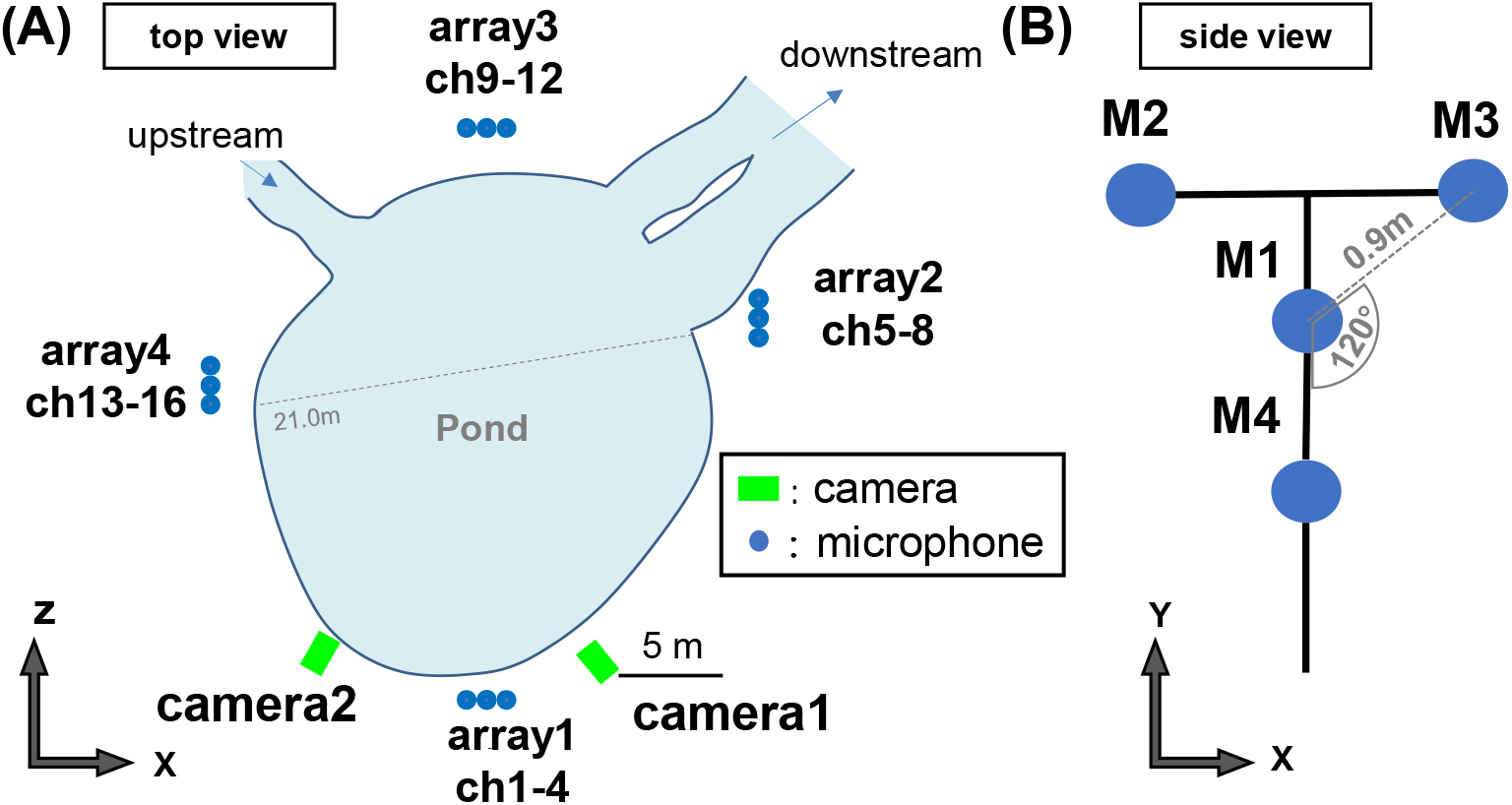
(A) Top view of the pond. A Y-shaped microphone array system was set up at four locations around the pond (total of 16 microphones) and two cameras were set up on one side of the pond. (B) Schematic of a Y-shaped microphone array system consisting of four microphones, with M1 at the center and three microphones (M2–M4) placed 0.9 m apart from each other.

**Table 1.** Overview of the meta-data and weather conditions.

| Day of recording | Sunset*<br>time | Recording<br>start | Recording<br>duration<br>[min] | Temperature<br>[°C] | Humidity<br>[%] | Weather |
| --- | --- | --- | --- | --- | --- | --- |
| 2018/6/18 | 19:14 | 20:00 | 130 | 13.4 | 79.7 | rain |
| 2018/6/19 | 19:15 | 20:15 | 110 | 14.0 | 79.7 | rain |
| 2019/6/23 | 19:16 | 20:27 | 23 | 15.0 | 77.3 | sunny |
| 2019/6/25 | 19:16 | 20:14 | 48 | 15.0 | 86.2 | cloudy |
| 2019/6/26 | 19:16 | 20:56 | 49 | 17.0 | 87.5 | sunny |
| 2019/6/27 | 19:16 | 20:29 | 61 | 17.4 | 83.4 | cloudy |

\* Data were from the National Astronomical Observatory of Japan.

Although this site is designated as a wildlife reserve by the local government, no permission was required for this survey because it did not include any endangered or protected species (Fukui et al., 2019), and we did not engage in any animal capture or habitat disturbance.

### Microphone array recordings

Bat echolocation calls were recorded using four T-shaped microphone array units, which comprised four omnidirectional electret condenser microphones (1/8-inch condenser microphones: models FG-23329-C05 and FG-23629-P16; Knowles Electronics, Itasca, Italy) (Fig. 2B, Fujioka et al., 2011). The distance between the central microphone, M1, and each of the three equally spaced outer microphones was 0.9 m (Fig. 2B). The four units in the microphone arrays (arrays 1–4) were arranged such that echolocation calls emitted from anywhere around the pond area could be recorded.

The echolocation signals recorded by the microphones were amplified and band-pass filtered (10∼250 kHz) using a custom-designed electronic circuit, and then digitized with 16-bit precision at a sampling rate of 500 kHz using a high-speed data acquisition card (PXIe-6358; National Instruments, Austin, TX, USA). The frequency response of the microphones was flat (±3 dB), and ranged between 10 and 250 kHz. The output signals were synchronously stored using a personal computer via a custom program created using LabVIEW 2011 (National Instruments). Recordings were saved as files every 10 minutes, and recording was stopped when the batteries ran out.

Sound data from the central microphone in each of the four microphone arrays (ch 1, 5, 9, 13) were analyzed. We used Cool Edit 2000 (Syntrillium Software Corporation, Phoenix, AZ, USA) to display the spectrograms of the sounds (128-point FFT, Han window with overlap for N-1) and extracted the terminal buzz signals with the clearest spectrograms for analysis. When the signal-to-noise ratio of a pulse was poor and extraction of the sound data was difficult, the data from the other channels were checked to determine the presence or absence of a terminal buzz. The post-buzz pause was calculated as the time between the last sound in the terminal buzz and the start of the next search pulse, as measured from the spectrogram images (Fig. 1A).

### Video recording and analysis

The foraging behavior of the bats at the pond was recorded using high-speed cameras (LT Recorder Pro (ver. 1.04; DITECT, Tokyo, Japan), in synchronization with the above-described sound recordings. The observation area was illuminated by infrared floodlights (LIR-CS88; IR LAB, Shenzhen, China) and the frame rate of the camera was set to 60 fps. An analog on/off control signal generated by a custom-made electrical circuit triggered video recordings, so that the video and sound data could be synchronously recorded and stored on the PC. The video recording was stopped when the hard drive of the computer was full (after approximately 30 minutes per measurement day).

Video images were analyzed visually using Dipp-Image Viewer (version 1.22; DITECT). In the first step, we identified scenes that showed a bat attacking prey, i.e., cases in which the tail membrane and hindfeet of a bat touched the water surface. These scenes were classified as “catch” or “failed”, and the “catch” group was further classified as “captured” (successful predation) or “dropped” (Fig. 3). Scenes were classified as “captured” when the bat caught the prey near the water surface with its feet or tail membrane and carried it away. Scenes where a bat caught prey but then dropped it were classified as “dropped”. The “failed” category contained scenes in which the presence of the prey on the water surface was confirmed after the bat had attacked.

**Figure 3.**
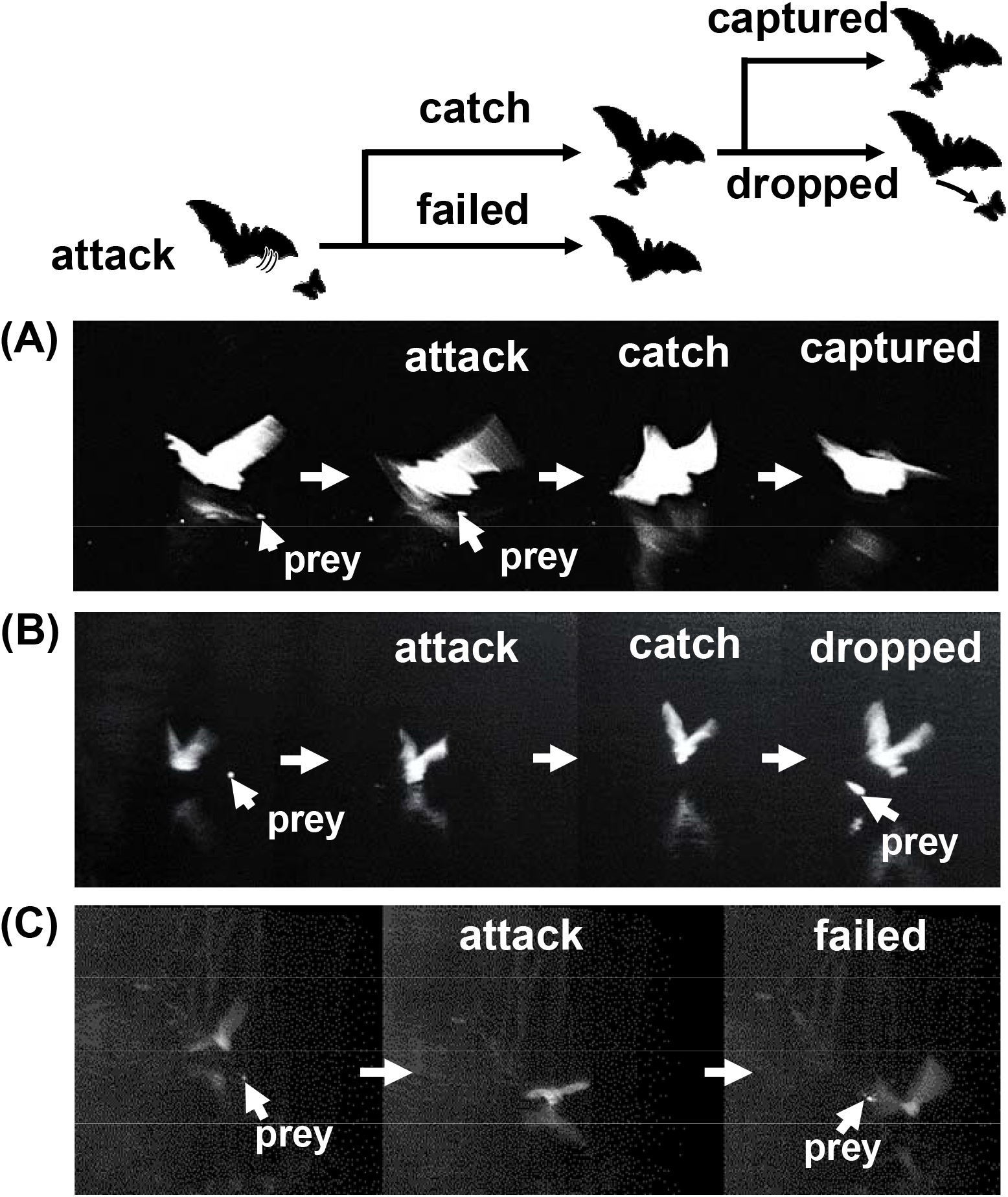
Images from video recordings of the three categories of attack outcomes. (A) “Captured”: the bat captures the prey near the water surface with its feet or tail membrane and carries it away. (B) “Dropped”: the prey is seen moving away from the bat after the bat initially caught it. (C) “Failed”: the prey can be seen on the surface of the water after the bat passed it (Movie 1).

### Statistical analysis

All statistical analyses were performed in the R environment for statistical computing (R Core Team 2020) and its extended packages. To test our hypothesis, generalized linear mixed models (GLMM) were built using the Template Model Builder package (glmmTMB_1.0.2.1; Brooks et al., 2017). Because several post-buzz pauses were measured in the same echolocation call sequence, and therefore from the same bat, we included “bat-ID” as a random effect in all models. Also, to account for any differences between recording events, we included the date of the recording as a random effect in all models. Hereby, the effect of bat-ID was nested within the date factor. The post-buzz pause was modeled as a function of the “attack outcome” factor (three levels; captured, dropped and failed). This variable was measured in milliseconds, and comprised integer values that could not have a value of 0. Therefore, we assumed a 0-truncated-Poisson distribution for all models.

To test for the potential influence of weather (three levels: sunny, cloudy and rainy) and nightly temperature, we included each variable in a separate model. We used the Akaike information criterion (AIC), corrected for small sample sizes, to select the best model (function model.sel, package MuMIn_1.43.17; Barton, 2020). We examined the quality of the model fit graphically using the functions in the DHARMa package (version 0.3.3.0; Hartig, 2020). We determined the overall model significance by using an χ^2^ test, comparing the best parsimonious model to its null model containing only the random effects (function anova, package stats_4.0.3, R Core Team, 2020). A χ^2^ _type-II-Wald_ test (function Anova, package car_3.0.10; Fox and Weisberg, 2018) was used to identify significant factors within the model, and Bonferroni correction was applied to all pairwise post-hoc comparisons between the levels of relevant factors (function lsmeans, package emmeans_1.5.3; Lenth, 2019).

## Results

We recorded echolocation sounds of the bat for a total of 421 minutes on six separate nights (June 18 and 19 in 2018, and June 23, 25, 26, and 27 in 2019). From this time period, 220 minutes of synchronized video and audio recording were collected and analyzed.

We classified a total of 137 attacks into the three categories (captured / dropped / failed) from the video recordings (Fig. 3, Movie 1). We found that bats kept hold of their prey in 51.8 % (“captured”, n = 71) of the attacks, and dropped their prey or failed to capture it in 29.2 % (n = 40) and 19 % (n = 26) of the attacks, respectively. From the 137 attacks confirmed via video, we identified 135 terminal buzzes. From these buzzes, 87 post-buzz pauses could be clearly measured from the spectrograms with a good signal-to-noise ratio (capture: n = 37, drop: n = 33, failed: n = 17). Based on the minimum AIC value, the model containing only the attack outcome factor was the best (Table 2). After graphically examining the model residuals, the fit was determined to be satisfactory. The best model explained significantly more variance than the null model (χ^2^ = 162.14, degrees of type-II-Wald freedom [df] = 2, P < 0.001), and the attack outcome factor was significant (χ^2^_type-II-Wald_ = 158.1, df = 2, P < 0.001). The post-buzz pause was longest in cases of successful capture, with a mean ± SE χ type-II-Wald value of 200 ± 11.14 ms (χ^2^_type-II-Wal_ test: capture vs. failed ratio = 0.573 ± 0.03, df = 82, P < 0.001, capture vs. drop ratio = 0.767 ± 0.03, df = 82. P < 0.001, Fig. 4). The post-buzz pause was shortest in cases of failed capture, with a mean value of 114 ± 7.05 ms (χ^2^_type-II-Wald_ test: drop vs. failed ratio = 1.340 ± 0.05, df = 82, P < 0.001), while the mean post-buzz pause in cases of dropped prey had an intermediate value of 153 ± 8.64 ms.

**Table 2.** Overview of models and their comparison based on the AICc.

| Model no. | Factors |  |  | Df | Loglik | AICc | Delta | Weight |
| --- | --- | --- | --- | --- | --- | --- | --- | --- |
|  | Capture outcome | Weather | Temperature |  |  |  |  |  |
| 1 | + |  |  | 5 | -885 | 1781 | 0 | 0.88 |
| 2 | + | + |  | 7 | -885 | 1785 | 4 | 0.12 |
| 3 | + |  | + | 19 | -874 | 1797 | 16 | <0.001 |
| 4 |  |  |  | 3 | -966 | 1938 | 158 | <0.001 |

**Figure 4.**
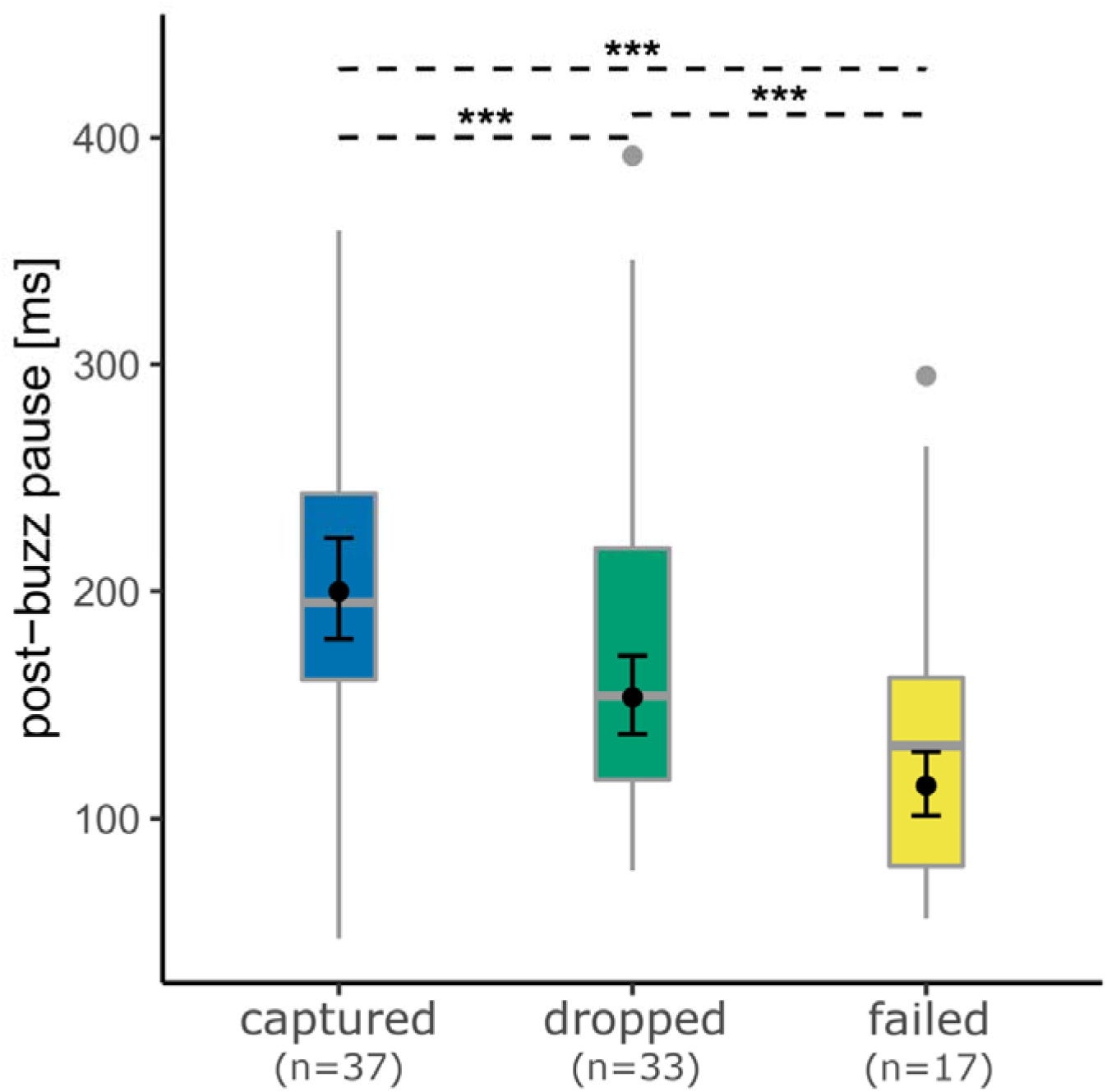
Post-buzz pause length for the various attack outcomes (captured, dropped and failed). Boxplots represent the raw data for each category. Means (filled circles) and 95% confidence intervals were derived for the best model (Table 2). Asterisks indicate significant differences between factor levels based on a Bonferroni-corrected post-hoc test conducted using the emmeans-package in R. ***p < 0.001.

We observed additional interesting behaviors in the video recordings. For example, the bat attacked small branches (“Attack branch” in Movie 2) or the same prey again after initially failing to capture it (“Repeated attack” in Movie 2). In addition, in cases of dropped prey, the bats usually dropped the item immediately after capture. However, there were a few cases where the bats dropped objects after holding them for a longer period (“Throw away” in Movie 2).

## Discussion

The terminal buzz emitted by bats has traditionally been treated as an acoustic indicator of an attack on a prey item (Britton and Jones, 1999; Griffin, 1958; Jakobsen and Surlykke, 2010; Racey and Swift, 1985). However, the presence of a terminal buzz is not sufficient to determine whether predation was successful. Various potentially indicative acoustic features have been examined, such as the lengths of terminal buzz I and II, the length of the post-buzz pause, and the average interpulse interval (IPI) after the terminal buzz (Acharya and Fenton, 1992; Britton and Jones, 1999; Surlykke et al., 2003; Übernickel et al., 2013). Laboratory experiments on foraging behavior have indicated that the presence or absence of predation affects the post-buzz pause (Acharya and Fenton, 1992; Britton and Jones, 1999; Surlykke et al., 2003; Übernickel et al., 2013). In contrast, no clear evidence for such a relationship could be derived from observations of wild bats during natural foraging (Britton and Jones, 1999; Surlykke et al., 2003). This was due to poor-quality acoustic data and the tendency for post-buzz pauses to be relatively short as a result of adaptation by the bats to the complex natural environment. To the best of our knowledge, this study is the first to show that the length of the post-buzz pause can be used to measure successful predation by wild bats during natural foraging.

### Prey selection during natural foraging in trawling bats

In the present study, *M. macrodactylus* bats dropped their prey during 30% of all recorded attacks (40 drops out of 137 attacks). Therefore, these bats do not appear highly capable of discriminating their prey, but rather make their prey selection after capture. This may explain the intermediate values of the post-buzz pause observed in this study.

Previous studies on prey selection in bats that predominantly hunt via trawling, like *M. macrodactylus*, have also suggested that bats have relatively weak target discrimination ability. Indeed, bats have been observed to sometimes attack objects instead of prey (Barclay and Brigham, 1994; Kalko and Schnitzler, 1989). For example, *Myotis lufigusus* and *M. yumanensis* did not appear to discriminate among targets, and attacked inedible targets (beetles and leaves) as well as edible prey of the same size (moths) during natural foraging (Barclay and Brigham, 1994). Other trawling *Myotis* species, such as *M. dasycneme, M. daubentonii*, and *M. capaccinii*, repeatedly attempted to capture inedible dummy targets placed on artificial surfaces that mimicked the reverberatory properties of water (Siemers et al., 2001). Therefore, this might represent a general prey selection behavior in trawling bats.

On the other hand, bats might drop not only inedible targets, but also edible prey unintentionally. For instance, in the “repeated attack” shown in Movie 2 the bat most likely dropped an edible prey item because it recaptured that item after dropping it. However, since this type of behavior was captured rarely by our cameras, there is a limitation to discriminate between these two types of behaviors in this study. In the future, knowing what the bats have caught would help distinguishing between drops of inedible or edible targets.

The post-buzz pause was also shown to be affected by prey size in a laboratory study with *Pipistrellus pygmaeus* (Surlykke et al., 2003). However, the authors reported that the prey size did not have a significant effect on the post-buzz pause in wild bats, according to acoustic recordings taken during natural foraging.

In summary, we found a clear relationship between the post-buzz pause and predation success in naturally foraging *M. macrodactylus*. However, further investigation regarding this relationship, including influencing factors such as prey type and size, will be needed to develop a reliable acoustic indicator of predation success in wild bats.

### Post-buzz pause applied: example of temporal changes in individual-specific predation success

To elucidate what insights could be gained by investigating post-buzz pauses during natural foraging, we recorded the foraging behavior of individual *M. macrodactylus* bats using four microphone arrays surrounding the pond, in the same experimental setting, for about 100 minutes starting at 20:04 on June 15, 2016. The recorded post-buzz pauses were analyzed based on their duration (represented by colors in Fig. 5) rather than whether predation was successful or not. During this recording period, a total of 70 bats visited the pond, 54 of which produced feeding buzzes (attacks) (note that the same bat might have entered the feeding area on multiple occasions). Bats that arrived during the first 30 minutes of the recording, which was the busiest period (20:30– 21:00), seemed to have shorter post-buzz pauses than those that arrived during the second 30-minute period. If predation success is related to the length of the post-buzz pause, then this observation suggests that predation success greatly varies depending on the time of foraging, and potentially also on the individual. This example shows that by using the post-buzz pause length as an acoustic indicator of predation success, we may be able to investigate detailed temporal aspects of foraging behavior in bats in feeding patches. Specifically, both the number of attacks and the length of the post-buzz pause can be used as indicators of predation efficiency during natural foraging.

**Figure 5.**
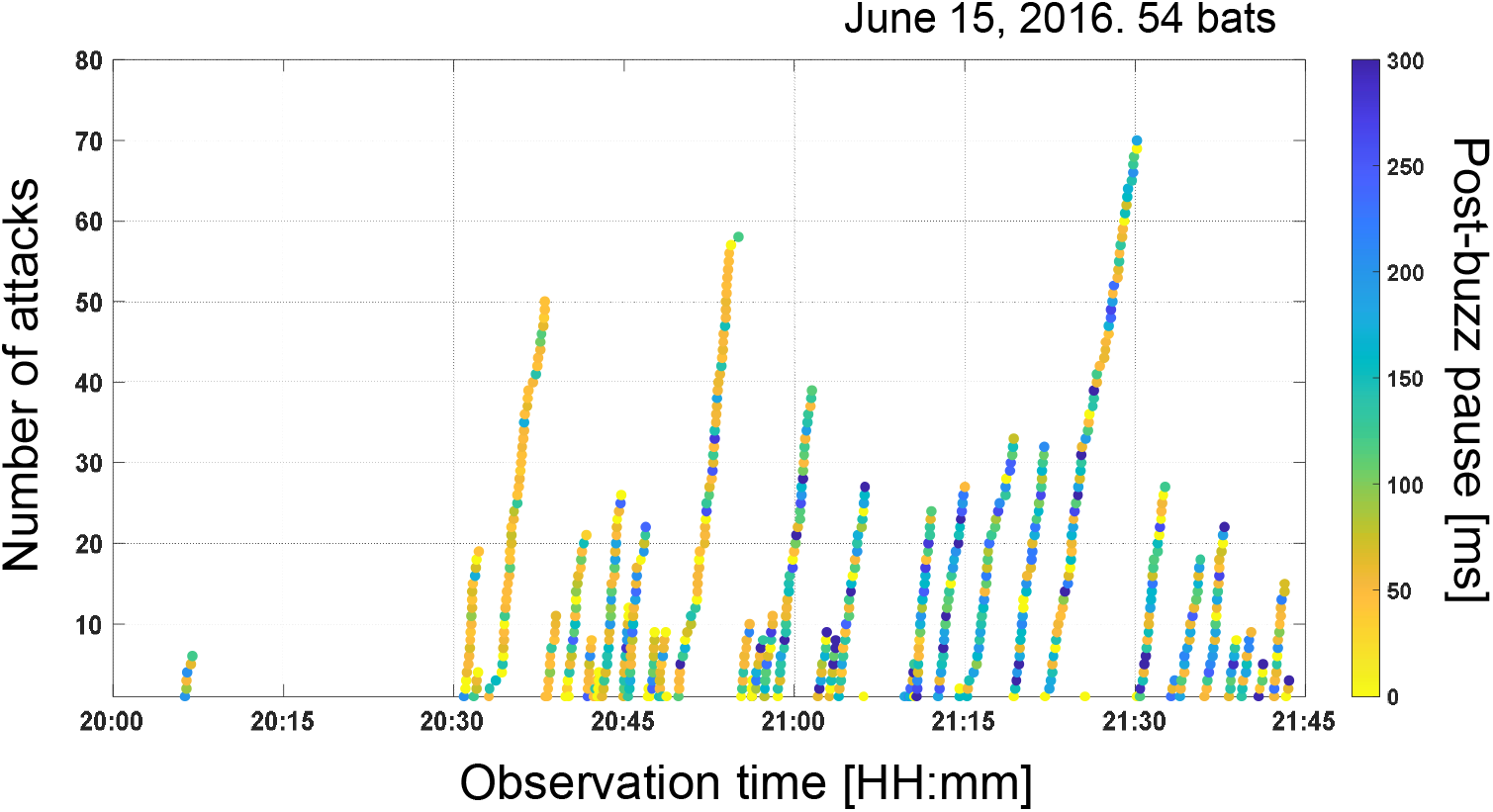
Temporal changes in the number of attacks by each bat that entered the foraging site on June 15, 2016. The time series shows the accumulation of attacks (filled circles) and the corresponding post-buzz pause lengths (color-coded). For reference, the average post-buzz pause was 200 ms for captured, 153 ms for dropped, and 114 ms for failed attempts.

## Conclusions

In this study, we conducted synchronized acoustic and video recordings of foraging behavior in wild Japanese large-footed bat *M. macrodactylus*. Our data showed that the bats either kept hold of their prey, dropped it, or failed to capture it after an attack. Overall, predation was successful in 51.8 % of attacks. Furthermore, the post-buzz pause was significantly longer in cases of successful predation than in the other two cases. This study is the first to identify differences in acoustic features between successful and unsuccessful predation in naturally foraging bats. Using the post-buzz pause as an indicator of successful predation will enable detailed investigations of wild bat foraging behavior, including foraging efficiency and, potentially, temporal changes in energy intake within feeding areas.

## Acknowledgments

We are grateful to Fumiya Hamai, Genki Nakai, Koki Asano, Mika Fukushiro, Yoshimasa Kitamura, Akihiro Nakae, Yoshiki Kubota, Yuta Sakuragi, Nozomi Sonobe, and Mika Kuroda for their assistance in conducting the field recordings.

## Data availability

The data supporting the findings of this study are available from the corresponding author upon reasonable request.

